# Advect, retain, sense: cilia-driven vortices deliver symbiont bacteria and flowborne cues to a mucociliary olfactory organ

**DOI:** 10.64898/2026.10.26.761908

**Authors:** Giancarlo Porcella, Janna C. Nawroth

## Abstract

Animals that acquire a microbial symbiont from the environment must find it against an overwhelming background of other bacteria. The bobtail squid *Euprymna* recruits its luminous symbiont *Vibrio fischeri* from seawater in which the symbiont occurs at fewer than ten cells per millilitre, and the host is transcriptionally primed for colonisation as soon as the first handful of symbionts become detectable at the light organ, too few to signal directly and implying a detection event upstream of it. We hypothesised that the paired olfactory organs, which sit at the entrance to the mantle cavity and hence in the path of all inhaled water, are that site. Here we characterise cilia-driven flow, particle capture and neuronal activity at the olfactory organ in hatchlings of *Euprymna scolopes* and *E. berryi* and in three non-symbiotic cuttlefish and squid species, i.e. species with no known light-organ symbiosis. In *Euprymna* the organ is a densely ciliated, mucus-coated epithelium packed with putative chemosensory cells and expressing the peptidoglycan-recognition receptor EsPGRP4, and its motile cilia drive a vortical flow that converges head-on onto the sensory face and persists to at least 96 h after hatching in colonised as well as aposymbiotic animals. Suspended beads accumulate there more strongly than at the light organ, and swimming symbionts follow the same flow yet are retained an order of magnitude more strongly than inert beads of the same size. Cells within the organ show spontaneous calcium transients, and the subset of cells and the firing rate change in response to waterborne stimuli, including live *V. fischeri*. In the three non-symbiotic species the equivalent organ sweeps water past its face rather than onto it, confining capture to the margins. Taken together, these findings suggest that in *Euprymna* the olfactory organ may concentrate, hold and detect arriving bacteria at a chemosensory surface before they reach the light organ.

**Significance:** Animals that acquire microbial partners from the environment must select them against an overwhelming background of other microbes. The bobtail squid, which recruits a single luminous symbiont from seawater, already exhibits colonization-specific changes in gene expression before meaningful numbers of symbiont cells have reached the target organ, suggesting a potential detection event upstream of it. Here we show that the squid’s paired olfactory organs, which lie in the path of all inhaled water and express receptors for bacterial wall components, are mucociliary sensory surfaces that draw water onto the organ’s center, where bacteria and bacteria-sized particles accumulate. Putative sensory cells in the *Euprymna* organ exhibit spontaneous calcium spikes and this activity rises with waterborne stimuli including bacteria, so the surface that captures the bacteria is also a surface that may sense them. In three non-symbiotic cephalopods the equivalent organ sweeps water past its surface and limits capture of flow-borne matter to the edges, suggesting a potential specialization of *Euprymna*’s organ in bacterial capture and detection compared to the other species. Mucociliary surfaces host microbial associations throughout the animal kingdom, and these findings suggest a general mechanism: motile cilia advect microbes to the surface, mucus immobilises them, and sensory cells embedded in that surface detect them.

## Introduction

Motile cilia are microtubule-based surface actuators that both propel and sense the extracellular fluid (1–3). On mucociliary epithelia they are classically described as clearance escalators, yet the same surfaces are the sites at which animals acquire and maintain their microbiota, and the mechanisms that reconcile clearance with selective engagement remain largely unresolved (4). In terrestrial vertebrates these tissues are internal and difficult to observe directly. Aquatic animals offer ciliated surfaces that can be imaged intact, in the living animal, while they are performing the function in question.

The binary association between species of the bobtail squid genus *Euprymna* and the luminous bacterium *Vibrio fischeri* is the most tractable of these systems (5). Most research has focused on *Euprymna scolopes* but recently developed laboratory culture of congener *E. berryi*, whose bacterial colonisation behaviours and host responses run largely parallel to those of *E. scolopes*, have enabled the study of successive generations in the laboratory and targeted gene knockout (6).

*Euprymna* hatchlings emerge sterile and must recruit *V. fischeri* from ambient seawater into a dedicated light organ, where the partnership is then maintained for life. The symbiont is extremely dilute in the ambient seawater: even within the squid’s own enriched habitat, *V. fischeri* is recoverable at no more than 900 colony-forming units per 100 ml, i.e., fewer than ten cells per millilitre, and is undetectable in water away from those habitats (7). Against a bacterioplankton background of about 5 × 10^5^ cells per millilitre on the continental shelf (8), the hatchling must find its partner at a dilution of order one in 10^5^. The sequence by which the symbiont colonises the light organ is stereotyped and experimentally accessible, and the molecular gates that follow initial contact at the light organ are well characterised, including mucus shedding, peptidoglycan recognition, nitric-oxide signalling, chemotaxis into the crypts (5, 9). Cilia contribute at the start of this sequence: ciliated fields on the light-organ surface generate flows that focus bacteria-sized particles, to the exclusion of larger ones, into sheltered zones adjacent to the entry pores (10).

The steps prior to this sequence, however, are not yet accounted for. Transcriptional changes consistent with preparation for colonisation occur across host tissues at a stage when only a handful of symbiont cells can be detected at the light organ, which is insufficient for direct biochemical host-microbe signaling (11). Moreover, light-organ ciliary beat frequency rises within fifteen minutes of exposure to environmentally realistic symbiont densities, which is before symbionts are physically attached (12).

Both observations imply that the host registers the presence of *V. fischeri* earlier, and somewhere other than at the light organ itself.

The paired olfactory organs are a candidate site. In coleoid cephalopods they sit on the head between the eye and the mantle opening, i.e., directly in the path of water inhaled into the mantle cavity, and carry densely packed ciliated and putative chemosensory cells whose axons project to a discrete olfactory lobe (13–16). Isolated squid olfactory receptor neurons respond to amino acids and other odorants (17), and behavioural work establishes distance chemoreception (18). A sensory surface in the inhalant path, wired to the central nervous system, could in principle register arriving bacteria on a faster timescale than biochemical signaling at the light organ.

Although ciliation is a common feature of cephalopod olfactory organs, whether those cilia are motile and what they transport has not been described. In other taxa, near-surface flow at an olfactory epithelium modulates chemoreception: in the zebrafish nose, motile-cilia-generated flow improves both the sensitivity and the temporal resolution of olfactory computation by controlling the rate at which odorant reaches and leaves the receptor surface (19).

Here we asked whether the cephalopod olfactory organ is a hydrodynamically active surface, and whether its activity differs between species that must capture a specific bacterium and species that need not. Specifically, we tested the prediction that follow from the hypothesis that the organ is specialised for detecting bacteria: in *Euprymna*, cilia-driven flow at the organ should promote the capture and immobilization of bacterial candidate particles near the sensory surface, whereas in non-symbiotic species, cilia should advect flow to the sensory surface without build-up of flow-borne matter.

## Results

### Bacteria-Sized Particles Accumulate Most Strongly at the Olfactory Organ

Suspended 1 µm fluorescent beads added to the water around an unsedated, i.e., normally ventilating, *E. scolopes* hatchling accumulated unevenly over external and internal body surfaces in a period of 5-10 minutes (Fig. 1 *a, b*). Relative to the surrounding water, bead signal increased approximately 7-11 fold at the olfactory organs, illustrating effective accumulation. At the light organs, the structure whose ciliated fields are known to recruit *V. fischeri* (10), the bead signal rose almost 3-fold. In contrast, the gill region, which serves as a control structure for unspecific capture from inhaled water, was only 20% brighter than background. On this whole-animal measure the olfactory organs are hence the strongest sites of particle accumulation on the hatchling.

**Figure 1.**
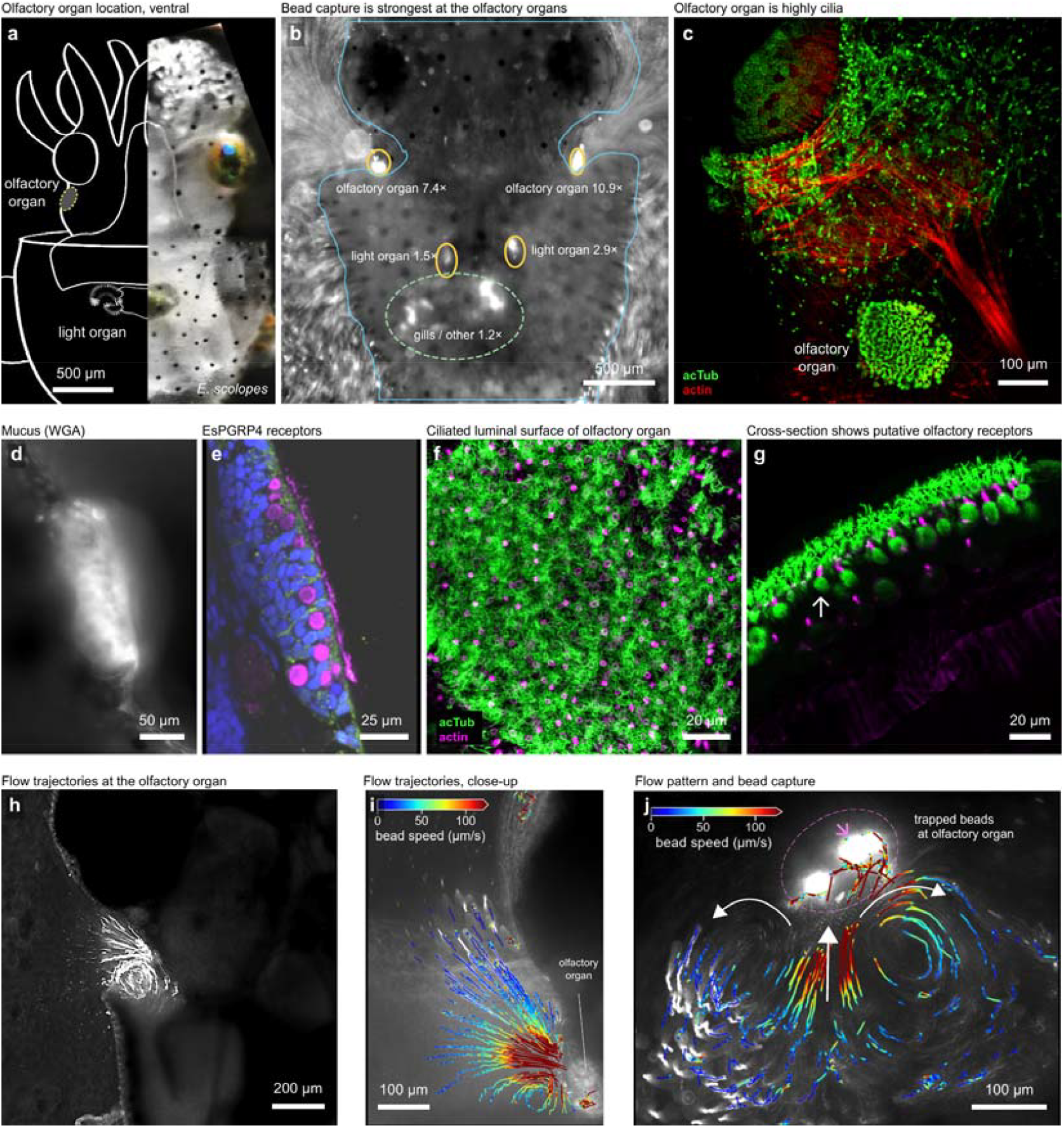
A ciliated olfactory organ in *Euprymna scolopes*. (**a**) Position of the olfactory organ seen from ventral side, light organ shown for reference. (**b**) Fluorescent-bead accumulation over the whole animal after approximately 5-10 min of exposure; bead signal relative to the surrounding water is indicated. Blue, body outline; orange, scored regions. (**c**) The organ in situ, below the eye; green, acetylated α-tubulin; red, F-actin. (**d**) Live wheat-germ agglutinin labelling of the organ (**e**) EsPGRP4 expression in the organ epithelium; blue, F-actin; green, EsPGRP4; magenta, α-tubulin. (**f, g**) Immunofluorescence of the ciliated face, top view and cross-section; acetylated α-tubulin, Alexa Fluor 488, green; F-actin (phalloidin Alexa Fluor 594), magenta. (**h**) Particle streaks at the olfactory organ in a live animal. (**i, j**) Bead trajectories at the organ in two fields of view, coloured by instantaneous speed. Scale bars: 500 µm (**b**), 200 µm (**h**), 100 µm (**c, i, j**), 50 µm (d), 25 µm (**e**), 20 µm (**f, g**). All panels are *E. scolopes*.

### The Organ Is a Ciliated, Mucus-Coated, Receptor-Bearing Epithelium

Immunofluorescence of the organ face resolved a dense apical ciliary carpet on top of a layer of acetylated α-tubulin-rich cells featuring actin tufts, consistent with cilia-containing chemosensory olfactory cells described in other cephalopod species (13) (Fig. 1 *c, f, g*). Live imaging of the ciliated surface revealed that the apical cilia are motile, with a ciliary beat frequency of ca. 5 Hz (see Fig. 6 *a*). The organ’s surface was labelled by wheat-germ agglutinin, consistent with a secreted glycoconjugate, or mucus, coat (Fig. 1*d*). The organ epithelium expressed Euprymna scolopes Peptidoglycan Recognition Protein (EsPGRP4) (Fig. 1 *e*), which was previously described at the light organ. There, exposure to bacterial peptidoglycan, a bacterial wall components released by ambient bacterioplankton in general, not by the symbiont specifically, causes the ciliated epithelium to release dense mucin stores within one to two hours of hatching, forming the substrate on which *V. fischeri* subsequently aggregates (20). EsPGRP4 is the receptor implicated in that response: it is predicted to act as a membrane receptor, it localises to those same superficial ciliated fields, and pre-incubating the tissue with an EsPGRP4 antibody reduces both mucus production and colonisation (21). While the present study did not test this mechanism, the presence of EsPGRP4 on the olfactory organ raises the possibility that this surface can also detect peptidoglycan and respond to it as the light organ does, by secreting mucus.

Hence, the olfactory organ combines in one surface three features that could facilitate bacterial encounter and detection: motile cilia, a mucus coat, and a receptor for a conserved bacterial cell-wall component.

### Cilia Drive a Vortical Flow That Converges on the Organ

In live, lightly sedated animals (i.e., with minimal mantel-driven ventilation flow), beads suspended in the surrounding water were advected along organised vortical paths focusing on the olfactory organ and exiting laterally (Fig. 1*h*–*j*). Time-averaged velocity fields in *E. berryi* resolved the same organisation: a vortex adjacent to the organ with streamlines converging on the central organ face (Fig. 2 *a, b*), and beads retained on the face itself (Fig. 2*d*). Near-surface speeds, measured from tracked particles within 30 µm of the surface, ranged from 98 to 261 µm s^−1^ across three *E. berryi* organs.

**Figure 2.**
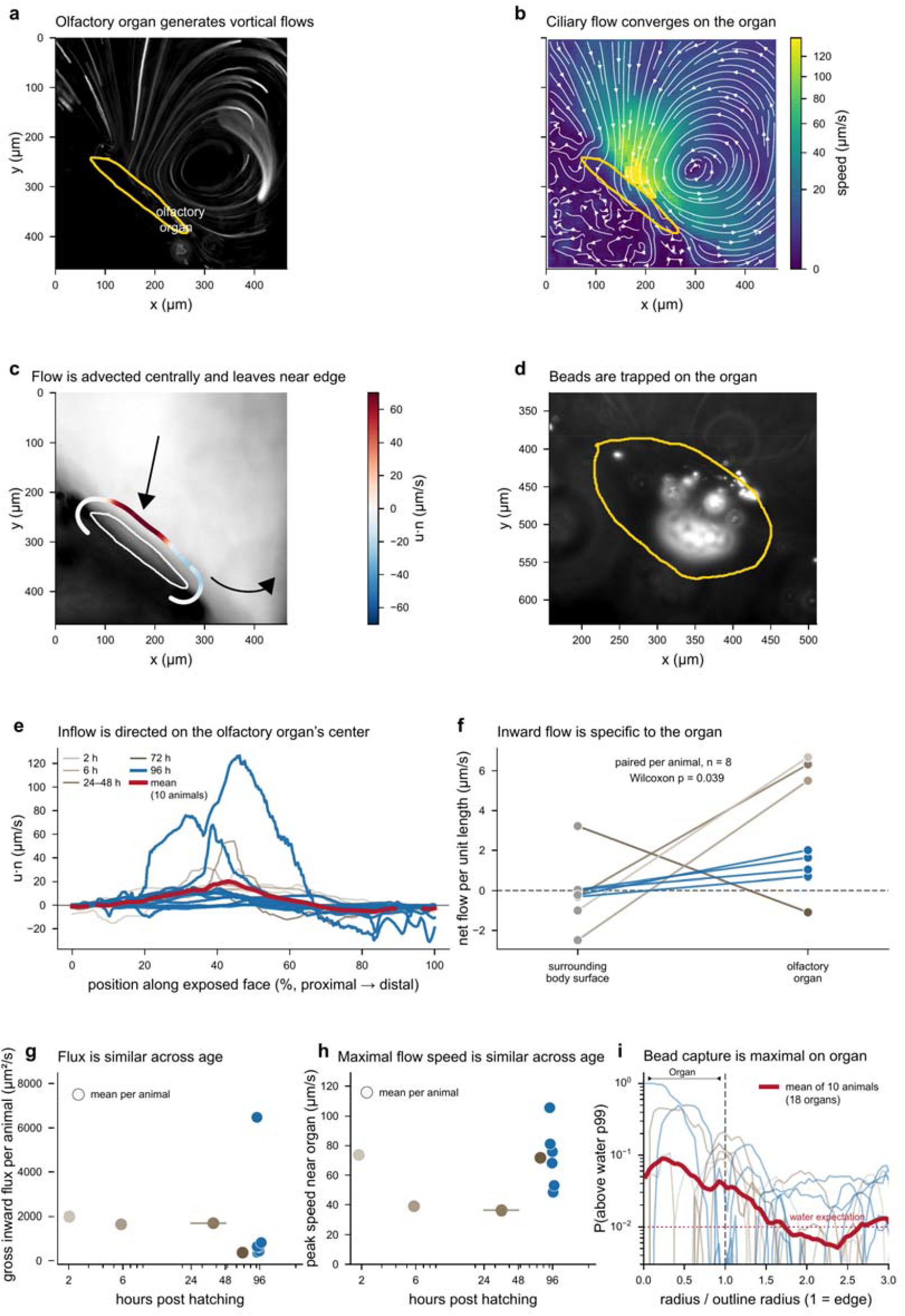
Cilia-driven flow at the *E. berryi* olfactory organ captures particles. (**a**) Bead streaks around the organ (96 h, left organ); yellow, organ outline. (**b**) Time-averaged velocity field with streamlines, same recording. (**c**) The arc at d/R = 0.60 on which flux is measured, coloured by the normal velocity component. (**d**) Beads retained on the organ (96 h, left organ). (**e**) Inflow profile along the exposed face for 17 organs, coloured by age, with the 10-animal mean in red. Each organ’s profile is normalised to its own largest excursion; absolute velocities span a 109-fold range across organs (1.2– 126.7 µm s^−1^) and are given in Figures 5*m* and 5. Six of the seventeen organs have their largest excursion in outflow and therefore reach −100% rather than +100%. (**f**) Net flow per unit length at the organ against a control stretch of body surface in the same recording, paired per animal (n = 8, Wilcoxon P = 0.039). (**g, h**) Gross inward flux and near-surface peak speed per animal against age; the 24–48 h animal is drawn as a bar spanning its range. (**i**) Probability of bead signal exceeding the 99th percentile of nearby water, against radius normalised to the outline (18 organs, 10 animals); dashed line, outline edge; dotted line, water expectation. One organ is excluded from **e** for a mis-registered ROI, which is why **e** reports 17 organs and **i** reports 18.

### Cilia-Driven Flow Is Directed onto the Organ in E. berryi

Across 18 organs from 10 animals spanning 2 to 96 h post hatching (11 organs from six animals at 96 h and seven organs from four animals between 2 and 72 h), flow measured on an open arc offset from the organ face was net inward in 14 organs (median net flux +610 µm^2^ s^−1^ across all 18 organs). The inward flux component peaked near the center of the organ’s face (Fig. 2*e*), along with mean speed normal to the surface (Fig. 5*m*). This spatial pattern was remarkably consistent while detected flow speeds spanned a wide range across organs (1.2–126.7 µm s^−1^).

Inward flow was specific to the olfactory organ rather than of the exposed body surface in general. In each recording in which both could be measured, net flow per unit length at the organ exceeded that at an organ-free stretch of body surface in the same field (n = 8 animals, Wilcoxon signed-rank P = 0.039; Fig. 2*f*).

### Flow at the Organ Persists to 96 h in E. berryi

The same head-on flow pattern was present between 2h and 96 h post hatching but three flow magnitude measures fell roughly two-fold with age, though no statistical significance was reached. The net flux of the eleven organs recorded at 96 h post hatching had a median of +560 µm^2^ s^−1^ compared to a median of +1477 µm^2^ s^−1^ at the 7 organs from earlier timepoints; (Mann–Whitney U = 45, P = 0.596). Similarly, gross flux showed a downward trend with age across the series (Spearman ρ = −0.66, P = 0.11; Fig. 2*g*). Mean flow speed along the measurement arc trended downwards with animal age (10.5 µm s^−1^ before 96 h; 5.6 µm s^−1^ at 96 h; U = 60, P = 0.056), while the peak speed near the organ remained stable (Fig. 2*h*).

Of note, the 96 h animals included both symbiotic and aposymbiotic individuals, which is important for interpreting the persistence of cilia-driven flow. At the light organ, successful colonisation both suppresses the ciliated recruitment apparatus and triggers its degradation. Mucus shedding from the superficial ciliated epithelium ceases within 48 h of colonisation, and the ciliated epithelial appendages begin to regress within hours of exposure to symbiosis-competent *V. fischeri* and have regressed completely by five days (22), through apoptosis triggered by the symbiont-released peptidoglycan fragment tracheal cytotoxin (23). If the olfactory organ were an equivalent structure, flow at it would be expected to decline once the animal is colonised. We did not observe such a decline. The two symbiotic animals gave animal-mean gross fluxes of 744 and 6646 µm^2^ s^−1^, against a range of roughly 500 to 1000 µm^2^ s^−1^ in the four aposymbiotic animals; neither symbiotic animal falls below the aposymbiotic range. With two animals against four no statistical comparison is possible so we report this as an absence of evidence for regression.

### Differential Retention of Live Bacteria and Inert Beads

A purely hydrodynamic account of capture predicts that a live bacterium and an inert particle of the same size, carried in the same flow, should arrive at the surface at the same rate and distribute identically around it: at these scales the capture mechanisms available to a surface in flow, i.e., interception, impaction, gravitational deposition and diffusion, all act on particle size and density rather than on whether the particle is alive (24). However, free-living *V. fischeri* propel themselves at approximately 66 µm s^−1^ (25), which is on the order of the cilia-driven flows we measured in the olfactory organs. Hence, the transport and fate of live bacteria may be different from that of passive bead. We tested this on a single organ, seeded first with 1 µm beads and then with GFP-labelled *V. fischeri* (at log phase) imaged with the same detector and analysed in the same distance bins (Fig. 3).

**Figure 3.**
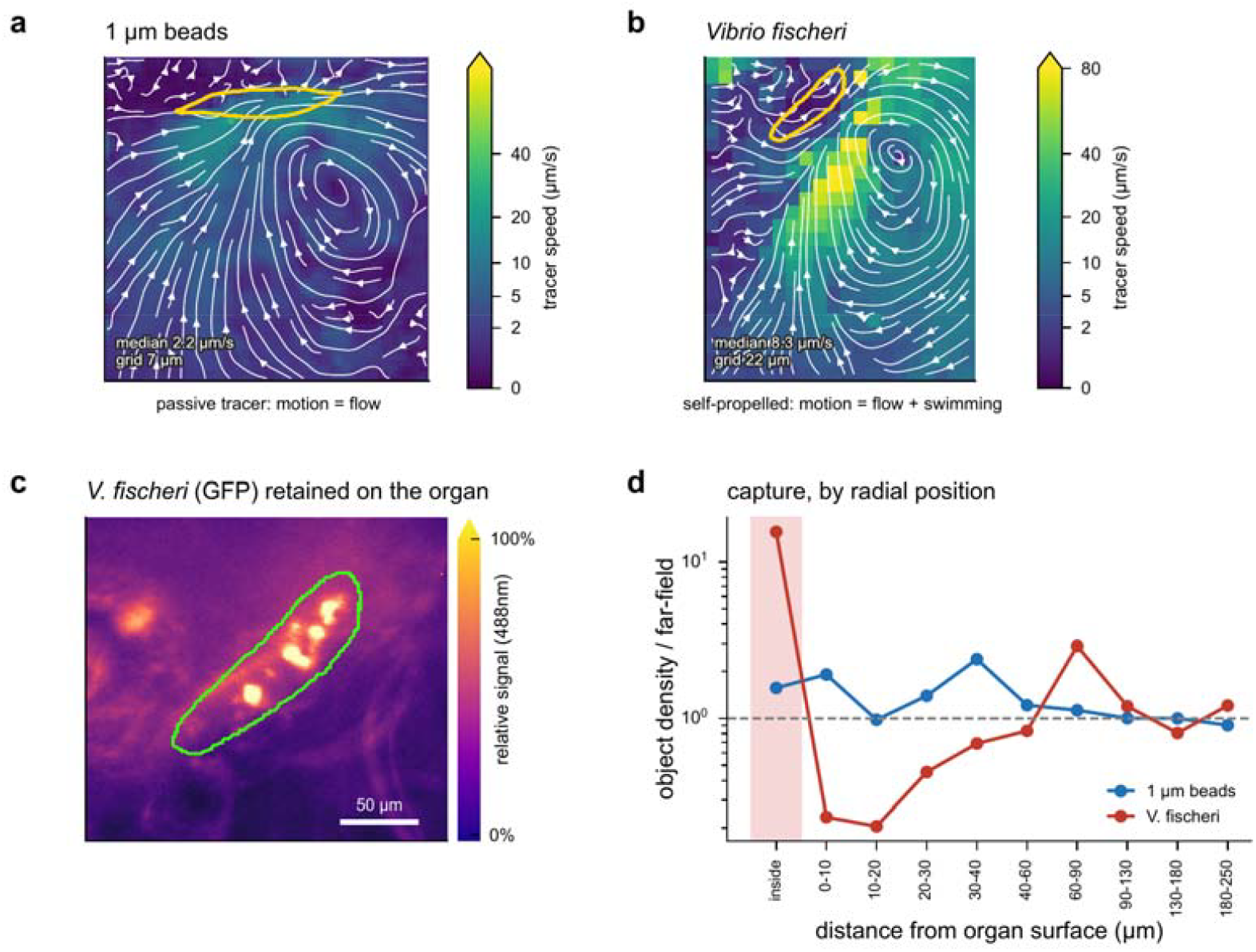
The organ handles passive beads and live *Vibrio fischeri* differently. (**a, b**) Time-averaged velocity fields at the same organ (96 h, right organ), seeded with 1 µm beads and with GFP-labelled *V. fischeri*; shared power-scaled colour. Bead motion reports flow alone (median 2.2 µm s^−1^); bacterial motion combines flow and swimming (median 8.3 µm s^−1^). Yellow, organ outline. (**c**) Time-average of the 488 channel, showing bacteria retained on the organ surface; green, outline. Scale bar 50 µm. (**d**) Object density relative to the far field by distance from the organ surface. Bacteria are 15.6× enriched on the surface itself yet depleted to 0.23× and 0.20× in the first two 10 µm shells of water, recovering to ∼1 beyond 90 µm. Beads show no such shell (1.6× on the surface, 0.9–2.4× at every distance).

The flow fields were comparable in organization but showed differences consistent with active propulsion in the bacteria: Flow-driven bead motion was slower (median 2.2 µm s^−1^) than the bacterial motion, which combines flow and active swimming (median 8.3 µm s^−1^). Both tracers were enriched on the organ itself, consistent with the bead accumulation in Fig. 2*i*: beads reached 1.6-fold the far-field density inside the organ outline and *V. fischeri* 15.6-fold, a tenfold difference between them. The two differed more sharply in the water just outside the surface, where *V. fischeri* was depleted to 0.23- and 0.20-fold in the first two 10 µm shells, recovering to approximately unity beyond 90 µm, whereas passive beads showed no such depletion shell and fluctuated between 0.9- and 2.4-fold at every distance (Fig. 3*d*).

The two tracers therefore gave opposite radial profiles at the same organ in the same flow. Depletion of bacteria from the water immediately adjacent to the organ, together with their tenfold-greater enrichment on it, is consistent with efficient removal of bacteria from the flow at the surface.

### Putative Neuronal Cells in the Olfactory Organ Are Spontaneously Active and Respond to Waterborne Stimuli

To ask whether the receptor layer of the organ is functionally active, we loaded 12 h aposymbiotic *E. berryi* hatchlings with the calcium indicator CaL520 AM and imaged a single plane of the organ face by light-sheet microscopy, acquiring the indicator and an autofluorescent reference channel simultaneously. The calcium dye preferentially accumulated in a subset of cells in the organ whose morphology and location beneath the apical ciliated cells was consistent with that of olfactory receptor neurons reported in other squid species (26) (Fig. 4 *a, b*). Though the animal was embedded in agarose during imaging, small movements and drift still occurred and hence each recording was registered on the reference channel to exclude motion artifacts (Methods). Loaded cells showed spontaneous, repetitive calcium transients (Fig. 4*c*), recurring at a dominant frequency of 0.10–0.71 Hz (median 0.23 Hz), with individual cells sustaining regular firing for the length of a recording. Active cells measured 2–79 µm^2^ in segmented area (median 25 µm^2^) and occupied a patch rather than the whole organ face, with neighbouring cells 22 to 62 µm apart, i.e., four to twelve cell diameters, and an arrangement within that patch indistinguishable from random placement (Clark–Evans *R* = 1.12). We verified the specificity of the signal by measuring unstained samples (Fig. S1). To our knowledge this is the first recording of calcium activity in the intact cephalopod olfactory organ; earlier calcium measurements in cephalopod olfactory receptor neurons were made in dissociated cells (27, 28).

**Figure 4.**
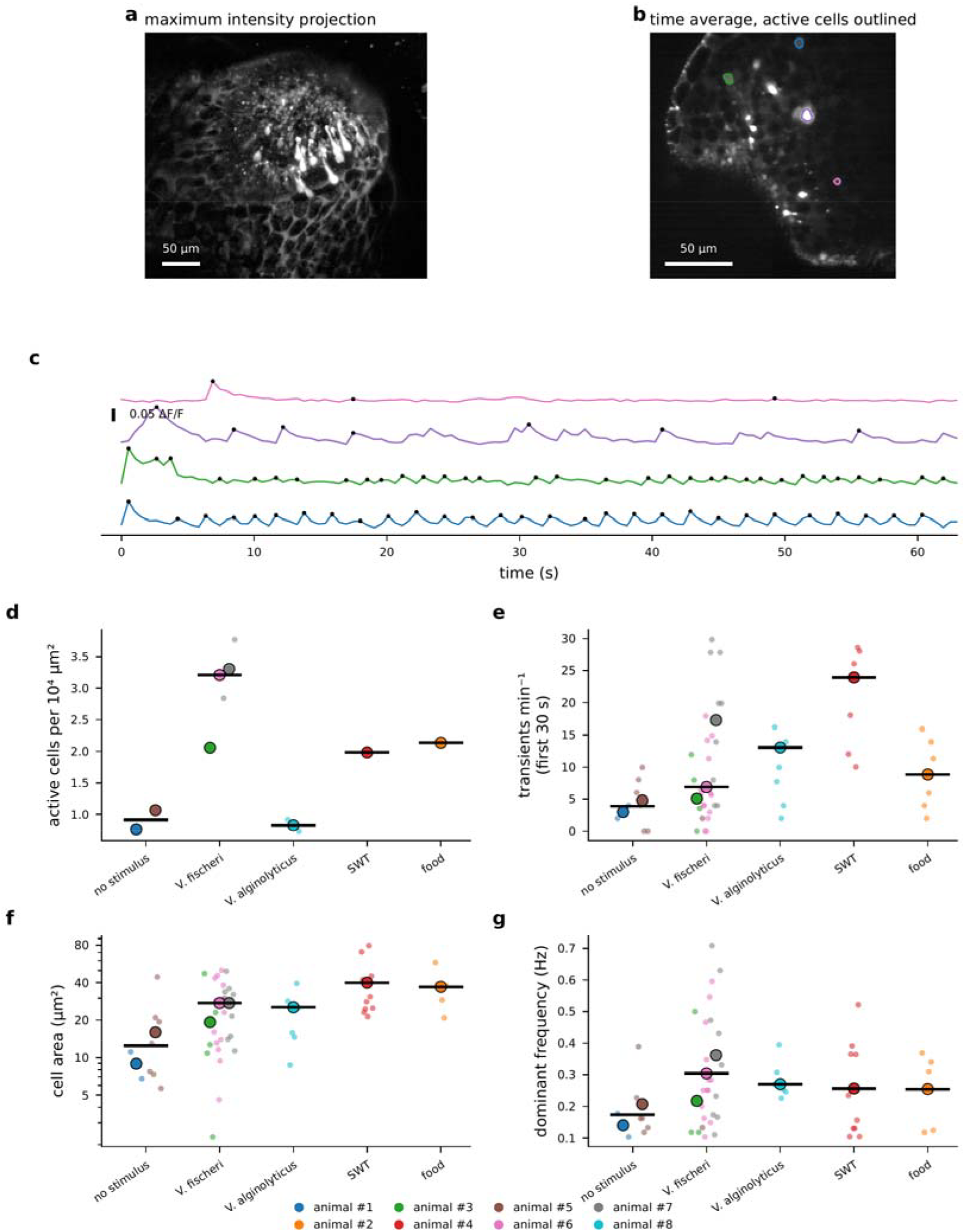
Calcium activity in the *Euprymna berryi* olfactory organ. (**a**) Maximum intensity projection of a z-stack through a CaL520 AM-loaded olfactory organ (0.46 µm px^−1^, separate animal); local contrast enhanced (CLAHE) to show stained cell bodies against the surrounding tissue. (**b**) Time-averaged calcium channel of one field from a *V. fischeri*-exposed organ; coloured outlines, the four segmented active cells retained after the artefact rules. (**c**) ΔF/F of those four cells, baseline removed; dots mark detected transients. (**d**) Active-cell density. (**e**) Transients per minute per cell, measured in a 30 s window common to all recordings. (**f**) Segmented cell area. (**g**) Dominant oscillation frequency (0.1–0.9 Hz band). In **d**–**g**, large circles are animal means and bars the median of those means; small points are the individual measurements, organs in **d** and cells in **e**–**g**, coloured by animal throughout. Quantities are aggregated cells to recording to organ to animal, so an organ imaged twice contributes once. Panels **d**–**g** use 10 organs from 8 animals, 60 cells across 14 recordings, after the three artefact rules; one recording is excluded as non-comparable. Scale bars (**a, b**) 50 µm.

Next, we tested the response of the organ’s cells to different waterborne cues, including symbiont *V. fischeri*, pathogen *V. alginolyticus*, crushed prey (mysid shrimp) and uninoculated growth medium (containing amino acids and other compounds known to activate cephalopod olfactory organs (17, (28))). Transient firing rate, average cell size of the responsive cell population, and dominant oscillation frequency each completely separated the stimulated from the non-stimulated group: every stimulated animal exceeded both unstimulated animals on all three metrics (6 animals against 2; one-sided Mann–Whitney U, *P* = 0.036) (Fig. 4*e*–*g*). Of note, since the stimulated pool included conditions containing none, or one of two different antibiotics, this comparison is not confounded by the selection agent carried in with the bacteria (Methods). The same structure appears in principal component analysis: over twelve per-cell metrics the first principal component carried 54% of the between-animal variance and the two unstimulated animals were the two lowest on it (Fig. S2).

### Active-Cell Density Is Highest in Organs Exposed to V. fischeri

The analysis did not distinguish one stimulus from another: condition identity carried no multivariate structure (silhouette coefficient −0.030), and the three *V. fischeri* animals lay no further from the vehicle animal than an arbitrary three-against-one split of the eight animals would place them (*P* = 0.39). What these data resolve is stimulated against unstimulated, not which stimulus. One measure nonetheless ordered the conditions consistently, and points to a potential sensitivity of the organ to *V. fischeri*. The number of active cells per unit area of organ face behaved differently from the per-cell measures, and was the one quantity that ordered the conditions (Fig. 4*d*). Organs exposed to *V. fischeri* gave a median of 3.21 cells per 10^4^ µm^2^ (four organs, n = 3 animals), against 0.91 in unstimulated conditions (two organs, n = 2 animals), a ratio of 3.5; crushed mysid shrimp gave 2.14 (one organ, n = 1 animal), uninoculated *V. fischeri* growth medium 1.98 (one organ, n = 1 animal), and *V. alginolyticus* 0.83 (two organs, n = 1 animal). The active-cell density of the four *V. fischeri*-exposed organs exceeded both unstimulated organs without overlap, but the single food-stimulated organ fell within their range, so active-cell density did not distinguish the symbiont from an unrelated chemical stimulus in this study. Besides being underpowered, the direct comparison is hampered by confounding factors such as differences in the antibiotic contained in the bacterial cultures (Methods). What these data support, however, is that the organ’s active-cell population is modulated by waterborne chemical stimuli and that the *V. fischeri* preparation produces the largest effect measured; what they do not establish yet is that the bacterium, rather than something delivered with it, is the cause.

### Cilia-driven Flow Patterns Limit Capture of Flow-Borne Matter In Non-Symbiotic Species

To investigate if *Euprymna*’s organ may have features specific to a symbiotic species, we recorded the equivalent organ and its cilia-driven flow in three non-symbiotic species: the cuttlefishes *Sepia bandensis* and *Metasepia pfefferi*, and the myopsid squid *Doryteuthis pealeii* (Fig. 5), revealing a clear difference between *Euprymna* and the non-symbiotic species. In *E. berryi* and *scolopes* water is drawn directly onto the organ’s center and expelled at its edges; in the other species the maximum inward flux is encountered at the organ’s edges, and water passes over or away from the organ rather than onto it (Fig. 5 *a-l*). In *E. berryi*, inward flux peaks near the center of the organ (median of 38% of the organ’s luminal length; n = 18, range 19–49%) whereas in the other species it peaks near the edge (median of 12% of organ length, n = 6, range 8– 20%; Mann–Whitney U = 106, P = 0.0001) (Fig. 5*m*). The two cuttlefishes place that inflow at opposite ends of the face: in *S. bandensis* it sits at the proximal end, near the eye (7%, 8% and 17% of face length), and in *M. pfefferi* at the distal end (79% and 87%), as it does in *D. pealeii* (88%). Which end water enters therefore does not generalise across the non-symbiotic species; what generalises is that it enters near an end at all, against the mid-face inflow of *E. berryi*.

The centrally-focused cilia-driven flow in *Euprymna* enriches the olfactory organ’s central surface with flow-borne bacteria-sized matter (Fig. 2 *e, i*), reflected in a higher average bead signal within the organ’s extent than outside of it (Fig. 5*n*). In contrast, the cilia-driven flow pattern in the non-symbiotic species tends to limit particle capture to the organ’s edge or outside of it. Together with the unfocused flow pattern, this is suggestive of a throughput, rather than capture, function.

**Figure 5.**
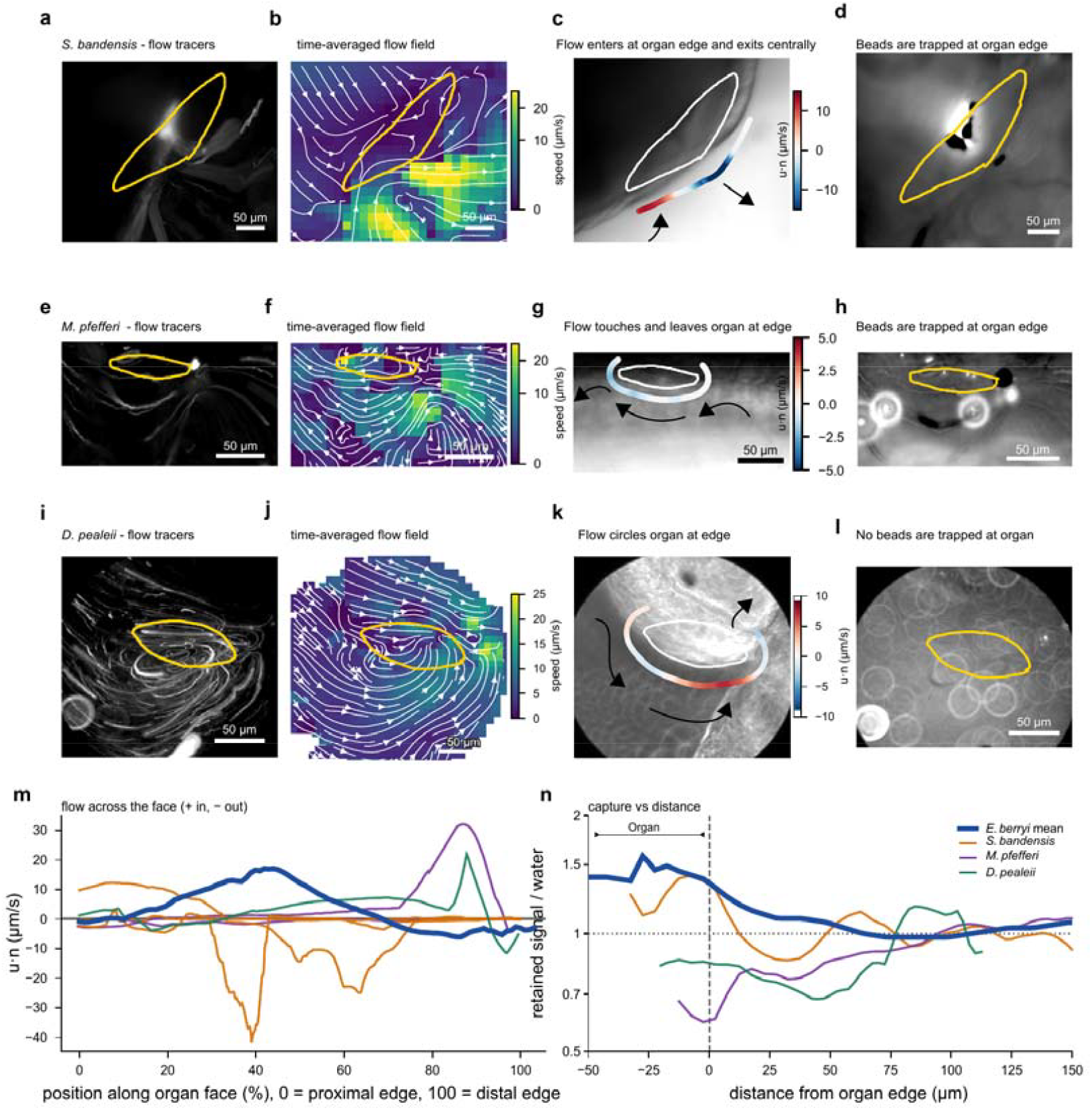
Flow pattern and bead capture differ between species. Rows: *S. bandensis* at 12 h (**a**–**d**), *M. pfefferi* at 48 h (**e**–**h**), *D. pealeii* at 3 h (**i**–**l**). The three are not age-matched; each is the recording of its species with usable bead streaks. (**a, e, i**) Bead streaks, i.e., a projection of the fluorescence stack; yellow, organ outline. (**b, f, j**) Time-averaged velocity field with streamlines, on a shared speed scale; the *D. pealeii* field in **j** is reconstructed from particle trajectories rather than by correlation, as described in the Methods. (**c, g, k**) The arc at d/R = 0.60 on which flux is measured, coloured by the normal velocity component; red is inward. Direction of surrounding flow is indicated by black arrows (**d, h, l**) Retained-bead maps of the same organs. (**m**) Normal velocity along the exposed face for all six organs of the three species, with the *E. berryi* mean of 10 animals in blue as a benchmark. The panel reports the shape, magnitude and proximal-distal orientation of the flow profiles. (**n**) Mean retained signal relative to each recording’s own 60–150 µm water band, against signed distance from the outline, with the *E. berryi* mean as a benchmark.

### Different Flow Patterns Correlate With Different Ciliation Patterns

The motile cilia are differently distributed over the organ in the tested species. Standard-deviation projections of phase-contrast recordings, in which moving structure writes a signal and stationary structure cancels, resolve ciliary motion across the entire organ face in *E. scolopes* at 24 h (Fig. 6*a*), matching the immunolabelled distribution (Fig. 1 *f, g*). In contrast, ciliary movement is confined to a band along the organ margin in the two cuttlefish *M. pfefferi* at 12 h and *S. bandensis* at 12 h (Fig. 6*b*–*c*), consistent with the tubulin-stained ciliation pattern reported for the cuttlefish *Sepia officinalis* (29). Pixel-intensity traces taken from the ciliated region are periodic in all three, at approximately 5 Hz in *E. scolopes*, 9 Hz in *M. pfefferi* and 6 Hz in *S. bandensis*. In *Euprymna*, a distinct point of phase reversal was observed roughly mid-way along the organ’s phase (Fig. 6*a*), consistent with the counter-rotating vortices observed above the organ (e.g., Fig. 1 *j*; *Figure S3)*. While ciliary beat was observed, no recording suitable for quantifying distribution or frequency was obtained from *D. pealeii*. Published staining data from *D. pealeii* paralarvae, the stage from which our recordings come, suggest that the olfactory epithelia are densely ciliated fields resembling those in *E. scolopes* (30). The absence of similar flow fields, however, suggests that ciliary beat organization differ between *D. pealeii and Euprymna*.

**Figure 6.**
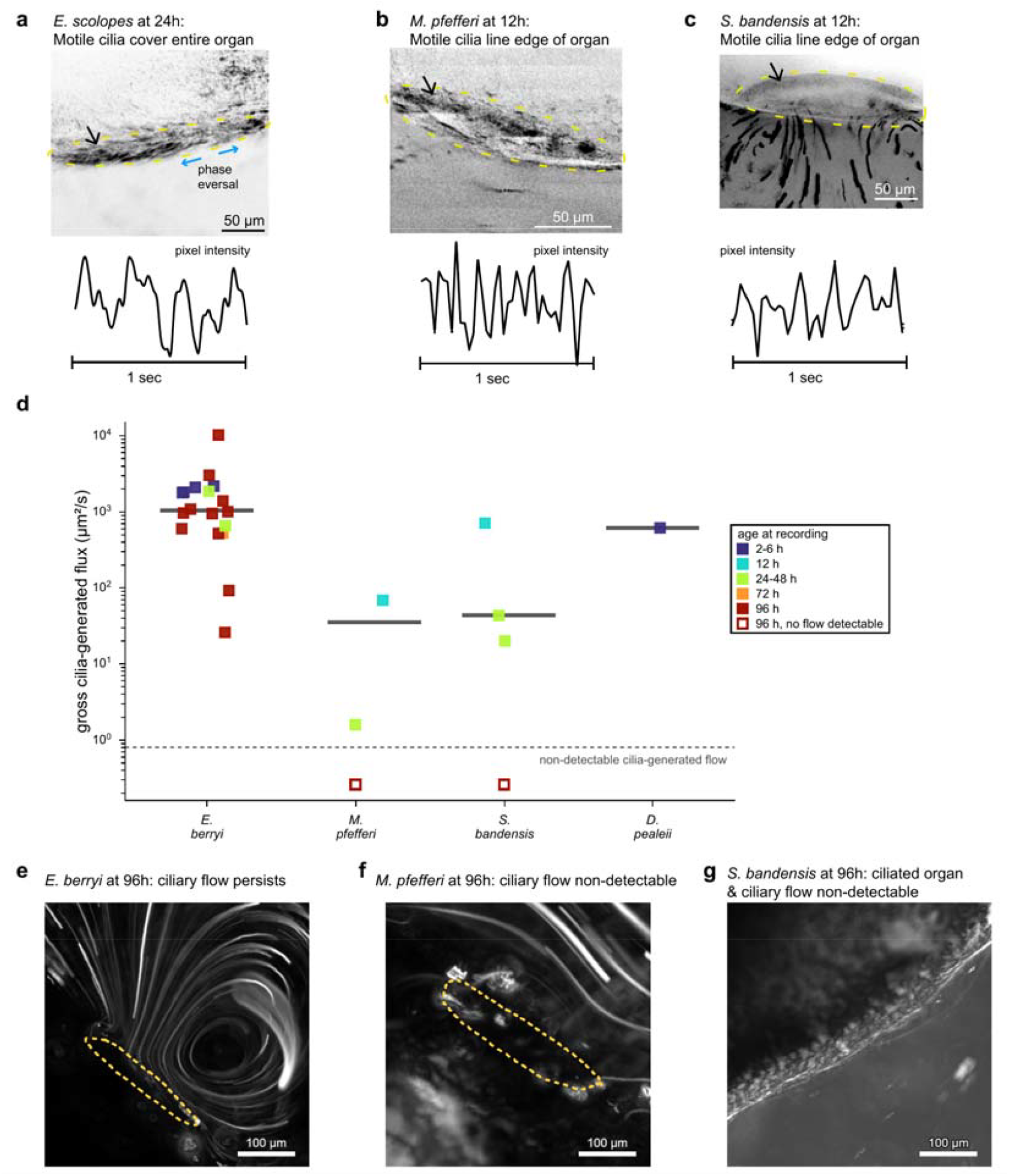
Ciliation and cilia-generated flux at the organ differ between species. (**a**– **c**) Distribution of motile cilia over the organ, from standard-deviation projections of phase-contrast recordings; the projection is inverted, so moving structure is dark and stationary structure cancels. Yellow, approximate organ outline. (**a**) *E. scolopes* at 24 h, with ciliary motion over the entire organ face and site of phase reversal is marked, with the opposing beat direction on either side indicated in blue. (**b**) *M. pfefferi* at 12 h and (**c**) *S. bandensis* at 12 h, with motion confined to a band along the organ margin. Beneath each image is a pixel-intensity trace against time taken from the ciliated region, revealing beat frequencies at approximately 5 Hz, 9 Hz and 6 Hz respectively. Scale bars 50 µm. No equivalent recording was obtained from *D. pealeii*. (**d**) Gross cilia-generated flux for every organ in the set, coloured by age at recording; bars, species medians over all ages. Open squares below the dotted rule are the two 96 h recordings in which no cilia-generated flow at the organ was detectable: *S. bandensis* retains no discrete organ to outline, and in *M. pfefferi* the motion in the field is driven by mantle movement rather than by cilia at the organ. Flux is measured on an open arc and is a gross magnitude at the exposed face; the direction of flow is shown in Figure 5**m**. (**e**–**g**) Representative recordings at 96 h, all shown as contrast-enhanced SD projections, so that moving tracer writes a bright streak while stationary structure cancels (**e** over its full stack; **f** and **g** over the first 200 frames of 388 and 387): *E. berryi*, with a vortex adjacent to the organ; *M. pfefferi*, with bead streaks crossing the organ outline and continuing past it; *S. bandensis*, a DIC stack in which neither a discrete organ nor ciliary flow is detectable. Dotted yellow, organ outline. Scale bars 100 µm.

### Cilia-Driven Flow Is Lower And May Disappear In Non-Symbiotic Species

Flow magnitude and its decline over time also separated the species. Restricted to ages at which flow in all four species were recordable (≤72 h), median gross flux was 1824 µm^2^ s^−1^ in *E. berryi* against 43.5, 35.2 and 617 µm^2^ s^−1^ in *S. bandensis, M. pfefferi* and *D. pealeii* (Mann–Whitney U = 39, P = 0.0082; Fig. 6*d*). At 96 h, *S. bandensis* and *M. pfefferi* had lost externally observable ciliation and the associated flow, in contrast to *Euprymna*, in which no such decline was observed.

## Discussion

We found that the cephalopod olfactory organ is a hydrodynamically active surface, and its activity differs between symbiotic compared to non-symbiotic species in ways consistent with the proposed role in symbiont detection: Organs in symbiotic species drive ambient fluid head-on to the mucosal surface, leading to apical capture of bacteria-sized particles in proximity to chemosensitive cells. Moreover, capture of free-swimming symbiont bacteria appeared to be especially effective compared to passive beads. In contrast, the organs in non-symbiotic species exhibited a variety of cilia-driven flow patterns that were circling or passing by the olfactory organ, and none of them focused flow directly onto the organ. As a consequence, trapping of bacteria-sized particles was limited to the organ edges or outside of the organ.

### Ciliation geometry accounts for the flow pattern

*Euprymna* carries motile cilia across the whole organ face, and the beat reverses direction half-way, associated with counter-rotating vortices drawing fluid to the center of the organ. In *S. bandensis* and *M. pfefferi* the moving cilia are confined to a band at the organ margin, a rim source, which sweeps fluid over the organ and out. Intriguingly, while *D. pealeii* may have densely ciliated fields similar to those in *E. scolopes*, the completely different flow field suggests that ciliary beat organization differs between *D. pealeii* and *Euprymna*. These species therefore present interesting model systems for studying the relation between ciliary beat organisation and flow transport function.

### Features of *Euprymna*’s olfactory organ support a role as symbiont detector

What distinguishes the symbiotic species’ organ is that water is directed onto the sensory face as a head-on stream *and* that material, including free-swimming bacteria-arriving there accumulates, likely aided by the mucus coating. This surface, which captures and retains bacteria, also expresses a receptor for bacterial peptidoglycan (EsPGRP4), consistent with a site at which arriving bacteria are detected. At the light organ, peptidoglycan from ambient bacterioplankton triggers the superficial ciliated epithelium to shed mucus, which is what *V. fischeri* then aggregates on (20), and EsPGRP4, the only member of the family predicted to act as a membrane receptor, is localised to those same ciliated fields and has been implicated in that response, since blocking it with antibody reduces both mucus production and colonisation (21).

The presence of that receptor on the olfactory organ therefore suggests a similar positive feedback: once the animal hatches, the ambient seawater carries peptidoglycan; peptidoglycan could be sensed by EsPGRP4; and mucus secreted in response would increase retention of bacteria, hence increasing the peptidoglycan signal. While each step is established at the light organ, the present study did not test it here at the olfactory organ yet.

### Two ciliated organs in one animal, serving related functions

At the light organ, two spatially segregated cilia populations act in sequence: long metachronal cilia focus bacteria-sized particles, to the exclusion of larger ones, into sheltered zones, where short randomly beating cilia mix a fluid environment containing host priming signals (10). The light organ sorts and delivers toward a destination, i.e., the pores leading to the colonisation crypts. Similarly, the olfactory organ draws water onto itself and retains material over its sensory epithelium. The same strategy, using ciliary beat to extract suspended material from a dilute background and accumulating it over a target site, therefore serves two related functions in the same hatchling, and the two organs share at least one molecular element in EsPGRP4.

### A putative “first detection” mechanism

The observation that motivated this work is that *Euprymna* is primed for colonisation before more than a handful of symbionts can be found at the light organ (11, 12). An external surface that concentrates and holds bacteria, sits in the inhalant path upstream of the light organ, and is innervated to a dedicated olfactory lobe, offers a route by which arrival could be registered early and relayed neuronally, far faster than chemical signalling from the light organ could achieve. The present data establish the hydrodynamic and retentive part of that proposal, and show that the receptor layer is spontaneously active and that its activity rises with stimulation with waterborne cues, including the symbiont. They do not yet establish the detection step itself. The data is insufficient to determine whether the organ distinguishes *V. fischeri* from other cues or from the medium it arrives in, or whether any signal triggers priming to colonization; what we provide are proof of concept data supportive of this role and a working preparation in which those mechanisms can now be explored.

### Reading neuronal activity at a beating ciliated surface

Mucociliary epithelia are difficult preparations for functional imaging: the surface moves, cells sit in different focal planes, and a z excursion brightens some cells while dimming others in the same frame, so the usual common-mode corrections cancel. Recording a reference channel alongside the indicator, registering on it, gating on per-cell focus and vetoing frames in which many structures step together addresses each of these. The same approach should transfer to other ciliated sensory and transporting epithelia, where motion has similarly limited what can be read out optically. Its main remaining limitation here is the two-dimensional nature of the recording that necessarily loses the information on the other planes. Volumetric acquisition, as described for other ciliated tissues (31) and adapted for simultaneous cilia, flow and calcium imaging could reveal the full mechanical and neuronal activity of the organ.

### Persistence beyond colonization

Flow at the organ is maintained at 96 h in *Euprymna*, long after initial colonisation is complete, while in the cuttlefishes it is gone, likely becoming an internalized structure (29). A surface used only for initial acquisition would not need to persist, as is the case in the ciliated light organ: mucus shedding stops within 48 h of colonisation (20) and the ciliated appendages bearing those fields undergo apoptosis beginning within hours of contact with symbiosis-competent *V. fischeri*, triggered by the bacterial peptidoglycan fragment tracheal cytotoxin acting together with lipopolysaccharide, and regression is complete within approximately five days (22, 23). Colonisation therefore both suppresses and removes the light organ’s recruitment machinery. The olfactory organ on the other hand remained intact and continued to drive flow at 96 h, flux and the head-on flow pattern persisted, and flow in the symbiotic animals lay within and above the aposymbiotic range. Therefore, the olfactory surface may persist and be used for continued sampling . There is a reason ongoing surveillance would be worth maintaining: The light organ expels most of its symbiont population at dawn each day, the epithelium remodelling and the residual bacteria then repopulating the organ over the following hours (32); that daily venting is itself what enriches the water *Euprymna* lives and breeds in, to as much as thirty times the *V. fischeri* density of comparable sites without squid (33). Detecting *V. fischeri* to find enriched breeding grounds, a prerequisite for successful colonization of offspring, would remain useful throughout life rather than only for the first hours of it.

### Cilia-driven fluid vortices are a common biological motif for delivering cargo to a surface

Cilia-generated vortical flows that manipulate flow-borne matter are not peculiar to *Euprymna’s olfactory* organ. At the *Euprymna* light organ, counter-rotating vortices generated by metachronal ciliary beat focus bacteria-sized particles into sheltered zones beside the entry pores (10). In reef-building corals, motile epidermal cilia covering the whole colony surface drive quasi-steady arrays of counter-rotating vortices that stir the water to 2 mm from the tissue and raise mass transfer across the diffusive boundary layer by up to 400% (34); that stirring redistributes photosynthetically produced oxygen laterally, away from the tissue in which it accumulates, and is thought to mitigate the resulting oxidative stress (35). In starfish larvae, external bands of motile cilia generate arrays of counter-rotating vortices, and by altering the number and position of those vortices the larva trades feeding against swimming, each behaviour having its own characteristic flow pattern (36). In the vertebrate brain, motile cilia on ependymal cells organise cerebrospinal fluid into a spatially structured flow network whose pattern switches periodically, subdividing the third ventricle into dynamically separated compartments (37, 38); modelling of the embryonic zebrafish ventricle, validated against in vivo transport measurements, indicates that the circulating ciliary flow governs the movement of large particles while diffusion dominates for small solutes (39).

What these systems share is not an evolutionary lineage but convergent functional solutions. A ciliated surface bounding a fluid volume, bordering walls, or containing phase-reversing sections generates recirculation rather than a unidirectional sweep, and recirculation raises both the residence time of material near that surface and the rate at which it is exchanged; bacteria fall in the size range that such flows transport effectively. Cilia-driven vortices hence appear to be a general solution in nature to control the transport and residence time of flow-borne matter near a tissue surface.

### Relevance beyond cephalopods

Horizontal acquisition of a microbiota at mucociliary surfaces is the general case, not specific to cephalopods, and the constraints are shared: a ciliated, mucus-covered epithelium must clear most of what arrives while engaging a specific minority. Mammals meet a version of the same problem with tuft cells, also called brush or solitary chemosensory cells, which are rare epithelial cells bearing a dense apical tuft of blunt microvilli over a tubulovesicular system rich in acetylated α-tubulin and DCLK1, distributed through the respiratory tract from the nasal cavity to the trachea and through the digestive tract from stomach to colon (31, 40, 41). They sense luminal contents using the transduction machinery of type II taste cells, such as α-gustducin, TRPM5, PLCβ2 and G-protein-coupled taste receptors including bitter receptors of the Tas2R family, and relay what they detect to immune and neuronal circuits (31, 42, 43). Described morphologically in the 1950s, their function was established only in 2016 (31, 42). The parallel to the cephalopod organ is that in both cases a chemosensory surface is embedded in a ciliated, mucus-producing epithelium at the interface with a microbially dense lumen, and meets it through a microtubule-rich apical specialisation. Whether cilia-generated flow contributes in the same way to microbial sampling at mammalian mucosal surfaces is, to our knowledge, untested.

### Limitations

Flux values are lower bounds: recordings were made at 10× magnification, which limits resolution close to the cilia, where velocity is highest. They are also limited to two dimensions, since the measurements are planar projections of a flow that is manifestly three-dimensional, so out-of-plane velocity, three-dimensional vortex structure and near-surface residence time are not recoverable; resolving them would require time-resolved volumetric imaging of flow and ciliary activity in the same preparation (38). Animals were lightly sedated and partially immobilised, which removes ventilation; the flows reported are the ciliary contribution in isolation, not the full hydrodynamic environment of a freely behaving hatchling. Sample sizes are small and uneven due to variable availability of hatchlings from different species at any given timepoint. Ciliary beat organization was only studied qualitatively and its relation to cilia-generated flow was not established quantitatively. Ciliary beat frequency estimates rest on measurements in one organ each and need to be confirmed. The identity of the active cells was not established: they were identified functionally, by calcium transients, and not by any molecular marker, so whether they are neuronal receptor cells is untested. In squid, olfactory receptor neurons transduce odorants through increases in intracellular Ca^2+^, and transduction occurs in their apical ciliary region (28, 44), which is consistent with the position and morphology of the cells recorded here but does not identify them. The calcium survey is underpowered to distinguish between the effect of individual conditions and confounding effects. Calcium imaging was confined to a single plane, so cells outside it and their activity are not represented.

### Next steps

Three classes of experiment follow directly. First, completing the comparative picture: later timepoints for *D. pealeii* and greater number of animals for both cuttlefish species across timepoints; immunofluorescence and live recording of ciliary organisation and flow, ideally in 3D, across age in all species; and EsPGRP4 staining in the non-symbiotic species, which would test whether the receptor tracks the capture phenotype. Second, testing the induction: whether peptidoglycan exposure elicits mucus shedding at the olfactory organ as it does at the light organ, and whether that changes capture. Third, testing systematically if the organ responds to *V. fischeri* specifically, for which the calcium preparation reported here provides the starting point. Subsequent studies could resolve whether ablating the organ or impairing its ciliary beat delays or reduces light-organ colonisation; and whether the organ, like the light organ, sorts flow-born matter by size and delivers different classes to different location on the surface, which could shape which receptor cells respond. A computational structure–function model relating ciliation geometry to the resulting flow would complete the quantitative understaning, and would allow studying a possible trade-off between capture/accumulation and sensing.

## Materials and Methods

### Animals

Hatchlings of *Euprymna berryi, Euprymna scolopes, Sepia bandensis, Metasepia pfefferi* and *Doryteuthis pealeii* were used, and age is reported as hours post hatching. All animals of all five species were obtained from the Marine Resources Center at the Marine Biological Laboratory, Woods Hole, MA, where *E. berryi* is cultured through its lifecycle (45). All husbandry and experimental protocols were in accordance with the EU 2010/63/EU and AAALAC guidelines for the use and care of cephalopods for research.

Egg clutches were hatched in 0.2 µm-filtered seawater supplemented 1:100 with penicillin–streptomycin. Aposymbiotic animals remained in this water and were never exposed to *Vibrio fischeri*. To create symbiotic animals, hatchlings were incubated with GFP-labelled *V. fischeri* at a final density of 10^5^ cfu ml^−1^ for 24h.

### Live imaging

Animals were sedated in 2% ethanol in artificial seawater and mounted ventral side up in a glass-bottom dish, with a coverslip used to restrain the animal without compressing it; on an upright stand the animal was inverted. The ventral aspect was imaged because the ciliated discs are exposed there and are covered by the mantle dorsally. All animals were sedated in this way except for the whole-animal bead-capture recording described below, which was imaged unsedated and therefore shows mantle movement.

Recordings were made on a Zeiss Axio Imager.M2 with an AxioCam 503 mono camera through a 1× adapter, running ZEN blue 3.3.89, in differential interference contrast and in epifluorescence, using an EC Plan-Neofluar 10×/0.30 Ph 1 (air) or a 40×/0.8 objective. Channels were brightfield, Alexa Fluor 488 and Alexa Fluor 594, with exposures of 1.3–150 ms depending on channel. With the camera’s 4.54 µm sensor pixels the image scale is 0.454 µm px^−1^ at 10× and 0.1135 µm px^−1^ at 40×; pixel size and frame interval were taken from the acquisition metadata of each file. The recordings analysed for flow have frame intervals of 22.6 ms or 25.5–25.6 ms.

Whole-animal bead-capture recordings were acquired on a Leica AF 6000 widefield stand with an HC PL FLUOTAR 5×/0.15 dry objective and a Leica DFC9000GTC sCMOS camera, 2048 × 2048 px over 2692.7 µm (1.3147 µm px^−1^), 1 ms exposure in the TXR filter cube, 896 frames at a 29.7 ms interval.

High-speed recordings of the ciliated face were acquired in Micro-Manager on a pco.edge 5.5 camera at 1 ms exposure, 800 frames at a 4.99 ms interval (200.4 frames per second).

### Tracers and bacteria

Flow was visualised with 1.0 µm carboxylate-modified polystyrene microspheres (FluoSpheres, Invitrogen cat. no. F8821, red fluorescent, 580/605 nm), diluted 1:1000 in filtered seawater, added directly before recording. For the whole-animal accumulation image (Figure 1*b*) the animal was left in the bead suspension for approximately 5-10 min before imaging. Bacterial experiments used *Vibrio fischeri* ES114, supplied by the laboratory of Spencer Nyholm as paired green- and red-fluorescent strains of the pES213-derived fluorescent-vector type developed for this species (46). The green strain expresses GFP constitutively and requires no selection; it was used for the flow and retention experiments. The red strain carries chloramphenicol resistance, was maintained at 5 µg ml^−1^ chloramphenicol, and was used for the calcium imaging. Both were supplied as colonies on seawater tryptone (SWT) together with liquid SWT of the same formulation. Cultures were inoculated from a fresh plate into liquid SWT, with chloramphenicol for the red strain only, and shaken at 28 °C for 4 h to mid-logarithmic phase. *Vibrio alginolyticus* was strain MD21-58, a genome-sequenced isolate from Stoney Beach, Woods Hole (2021), carrying pKJK5::GFP with kanamycin resistance. It was recovered from frozen stock into 3 ml Seawater Complete (SWC) medium with antibiotic added at 1:100 and grown for 12 h at 30 °C with shaking; working cultures were subsequently started from this culture and grown for 5.5 h under the same conditions to mid-logarithmic phase. Culture density was not measured by optical density but estimated from time in culture, following the conversions supplied by the source laboratories; on that basis mid-logarithmic cultures of both species were taken to contain approximately 10^8^ cfu ml^−1^. For acute imaging and for colonisation assays, bacteria were added directly at a final density of 10^5^ cfu ml^−1^. For calcium imaging, animals were exposed to approximately 10^6^ cfu ml^−1^ for 10–30 min.

### Immunofluorescence and lectin labelling

Whole animals were fixed in 4% paraformaldehyde in PBS overnight at 4 °C, then washed and permeabilised in PBS containing 0.1% Triton X-100. Samples were incubated overnight at room temperature with mouse monoclonal anti-acetylated tubulin (Sigma-Aldrich T6793) at 1:200, washed three times in PBS, then incubated for 2 h at room temperature with goat anti-mouse IgG (H+L) cross-adsorbed Alexa Fluor 488 (Invitrogen A-11001) at 1:500 together with Alexa Fluor 594 phalloidin (Invitrogen) at 1:500, protected from light. After three PBS washes, mantle tissue was dissected to expose or isolate the ciliated discs; and samples were mounted in SlowFade Glass antifade mountant (Invitrogen) in an ibidi glass-bottom dish under a No. 1.5 coverslip.

Stacks were acquired on a Leica TCS SP8 confocal with an HC PL APO CS2 40×/1.30 oil objective, scanning xyz at 700 Hz with 4-line averaging, a pinhole of 1 Airy unit (65.3 µm), and 2048 × 2048 px over a 153.2 µm field (0.0748 µm px^−1^). Cilia (Alexa Fluor 488) were excited at 488 nm and collected over 502–526 nm; F-actin (Alexa Fluor 594) was excited at 552 nm and collected over 590–613 nm.

For lectin labeling, live animals were incubated with wheat-germ agglutinin, Alexa Fluor 488 conjugate (Invitrogen W11261), at 3 µl ml^−1^ for 15 min, protected from light. Imaging was conducted in live, mildly sedated animals.

### EsPGRP4 labelling

EsPGRP4, a peptidoglycan-recognition protein, was detected with a rabbit antiserum raised against a synthetic peptide unique to it, in the laboratory of Margaret McFall-Ngai. The antiserum is not commercially available and is used as whole serum rather than purified IgG, at 1:100. The stain reports the presence and distribution of the receptor protein.

### Image projections

Maximum-intensity projections of fluorescence recordings render bead paths as continuous streaks, and standard-deviation projections emphasise moving objects against a static background; the projection used is identified in each figure legend. For DIC recordings a range projection (per-pixel maximum minus minimum) is used, which resolves moving particles against the bright field. Display ranges are percentile-based and stated per panel. Images used for measurement were not contrast-adjusted.

### Particle image velocimetry

Velocity fields were computed in PIVlab 3.14 (47) (MATLAB R2025a) from the fluorescence recordings by FFT cross-correlation with window deformation in two passes, a 64 px interrogation window at 32 px step followed by a 32 px window at 16 px step, giving 50% overlap and a final vector spacing of 7.3 µm. Sub-pixel displacement used a Gaussian estimator. Vectors were validated with a global standard-deviation filter at 8 SD and a local median filter at a threshold of 3, missing vectors were interpolated, and fields were smoothed with a parameter of 0.2. Spatial calibration used a 1 mm reference distance. A 3 px Wiener denoise was applied; image high-pass filtering and intensity capping were not used.

Each exported velocity field is a time average over the frame pairs of its recording rather than an instantaneous field, which is necessary at the seeding densities here: a single interrogation window contains well under one particle in any one frame pair, so instantaneous correlation returns noise while the average over many pairs is well conditioned.

Velocity was computed between frames separated by *N* frames, with *N* chosen per recording so that particle displacement exceeded the correlation noise floor. All *E. berryi* fields and the *S. bandensis* and *M. pfefferi* recordings at 12 h use consecutive frames; *S. bandensis* at 12 h uses *N* = 2, *S. bandensis* at 24–48 h *N* = 5 and *M. pfefferi* at 48 h *N* = 8. The *D. pealeii* field is not a correlation field; its equivalent quantity is the 60-frame lag (120 ms at 500 frames per second) over which particle displacements were measured. The effective intervals across the dataset are 22.6 ms (*E. berryi*), 51.2 and 113.1 ms (*S. bandensis*), 120.0 ms (*D. pealeii*) and 181.0 ms (*M. pfefferi*), so the *D. pealeii* interval sits inside the range and is shorter than the *M. pfefferi* one; what distinguishes that recording is its pixel size, 0.209 against 0.454 µm px^−1^, which resolves a flow 2.2× slower at matched pixel displacement and is why the long lag was needed to clear the correlation floor at all. Entered time steps were checked against the per-frame timestamps of each source stack and corrected where they disagreed.

Re-running the three recordings that required *N* > 1 across *N* = 1–20 changed the implied speed by at most 11% for the two slowest. For the fastest of them the estimate depends on *N* by a factor of 1.8, and the value reported is the one measured at the smallest usable separation, which is the larger of the two and therefore conservative with respect to the species difference.

### Doryteuthis pealeii acquisition and velocimetry

The *D. pealeii* recordings were made using a Photron high-speed camera at 500 frames per second, 1024 × 1024 px, 8-bit, with a measured pixel size of 0.2092 µm. Three stacks of the same field were acquired from one 3 h hatchling: two transmitted-light stacks and one fluorescence bead stack (5176 frames). The velocity field was reconstructed from particle trajectories. Beads were detected and linked in TrackMate 8.1.3 under Fiji on the fluorescence stack. Trajectories whose net displacement was below 5 µm were discarded as stationary debris. Velocity was taken as the displacement over a lag of 60 frames (120 ms). The samples were reduced to a field on a 6 µm grid by Gaussian-weighted local averaging (radius 12 µm, σ = 6 µm), with a node left undefined where fewer than 10 samples fell within its radius. The averaging radius was set to match the effective resolution of the 7.26 µm PIVlab fields rather than to maximise detail, so that the flux measures stay comparable across species.

### Organ outline and the control arc

Three regions were drawn in ImageJ for each recording: a closed freehand outline of the organ on the face-on projection, a freeline tracing the tissue border on the DIC projection, and a mark at the organ centroid.

Flux was evaluated on an open arc offset from the exposed face into the water by *d* = 0.60 *R*, fixed across the dataset. The arc is open because net flux through a closed contour surrounding an impermeable surface is zero by mass conservation, and the offset clears the two interrogation windows nearest the boundary, within which PIV is unreliable. An offset sweep from 0 to 80 µm is reported for every recording.

### Flux measures

With *u·n* the velocity along the inward normal and *ds* the arc element, gross inward flux is Q_in = ∫ max(u·n, 0) ds, net flux is Q_net = ∫ u·n ds, and their per-unit-length forms are q_in = Q_in / L and q = Q_net / L, with *L* the arc length carrying a finite velocity. On an open arc Q_net measures net through-flow: far-field water drawn into contact with the organ and flushed out through the open ends of the arc.

Because an open arc is not invariant to a uniform background current, the far-field velocity was measured for every recording as the median over grid nodes more than *R* + 150 µm from the outline centroid, and its contribution computed as Q_drift = U_x Σ n_x ds + U_y Σ n_y ds. Across the 23 recordings with a far field beyond that radius the drift has a median of 0.74 µm s^−1^ and a maximum of 5.86 µm s^−1^, and Q_drift is a median of 8% of Q_net; removing it changes the sign of two organs, both from values near zero, and does not change the species result. The *D. pealeii* field is 214 µm across and contains no node beyond *R* + 150 µm, so its drift was measured instead over nodes more than 80 µm from the outline, giving 2.92 µm s^−1^; subtracting it changes that organ’s net flux by 3% (+480 to +496 µm^2^ s^−1^).

### Particle tracking

Beads were detected and linked with custom Python code; the *D. pealeii* trajectories are the exception and come from TrackMate, as described above. Detection used a difference-of-Gaussians band-pass (σ = 1 and 4 px), thresholded at 6 median absolute deviations above the local background, located as local maxima with a minimum separation of 3 px, and refined to sub-pixel position with a 5 × 5 intensity-weighted centroid. Frames were linked pairwise by optimal assignment with a maximum displacement of 12 px, raised to 20 px for the fastest recording. Trajectories shown in figures are those of at least 20 frames with a straightness of at least 0.40; where a figure also required coverage of regions where the flow is fastest, trajectories of at least 6 frames with a net displacement of at least 10 µm and a straightness of at least 0.30 were added where they cross grid cells in the top decile of speed, and both criteria are stated in the legend. Only fluorescence recordings were tracked.

### Accumulation and capture

Bead and bacterial accumulation on the organ was measured from the time-averaged fluorescence image as the mean within the outline. A pixel was scored as carrying signal if it exceeded the brightness of the 99th percentile of water pixels within 200 µm outside the outline; the same statistic was computed at the 95th and 99.9th percentiles, and an organ was counted as accumulating only if it exceeded the water by the same factor at all three thresholds. Radial profiles are reported against radius normalised by the outline’s own radius at that angle. For the bead-versus-bacteria comparison, object density was counted per unit area in signed distance bands from the organ surface.

### Ciliary beat frequency

Ciliary beat frequency was measured in high-speed stacks by counting the average number of oscillations per second seen in kymographs derived from regions with visible ciliary beat.

### Calcium imaging

#### Dye loading and stimuli

CaL520 AM (AAT Bioquest 21130), 50 µg, was dissolved in 8 µl DMSO and added to 4 ml of 0.2 µm-filtered seawater (1:500). 12 h aposymbiotic hatchlings were incubated in the dye solution for 2–3 h (2 h for the 2023-08-30 cohort, 3 h for 2023-08-29), rinsed for 1 h in filtered seawater, then given the stimulus; no stimulus overlapped the loading period. Bacterial and vehicle stimuli were delivered by adding 40 µl of mid-logarithmic suspension to the 2 ml of seawater surrounding the animal (1:50, approximately 10^6^ cfu ml^−1^) and incubating 10–30 min. Bacteria were not added to an already-mounted animal, because free-swimming bacteria crossing the field interfere with transient detection; by the end of the incubation bacteria were visibly adherent to the organ, so the stimulus remained present during imaging, the rinse removing the free-swimming population and not the bound one. The vehicle was fresh, uninoculated chloramphenicol-containing SWT, the medium *V. fischeri* was grown in, delivered identically. For the food stimulus, mysid shrimp were crushed in filtered seawater and a droplet added immediately before mounting, remaining with the animal throughout the recording. Unstimulated and no-dye animals were held in filtered seawater during the stimulus phase. All animals were then sedated for 10 min in 2% ethanol in filtered seawater, embedded in 4% agarose and imaged at room temperature.

#### No-dye controls

Three animals received the same 1:500 DMSO without the indicator and the same sedation, embedding and acquisition settings; for one cohort a dye-loaded recording at identical settings is available as a matched pair. These were tiled with cell-sized regions and passed through the identical registration, ΔF/F and event-detection pipeline, giving a measured false-positive floor rather than an assumed one.

Autofluorescence accounts for approximately 80% of resting fluorescence, so reported ΔF/F understates the indicator’s own ΔF/F roughly fivefold.

#### Acquisition

Zeiss Lightsheet Z.1, SPIM mode, W Plan-Apochromat 10×/0.5 water immersion (n = 1.333), two pco.edge 4.2 sCMOS cameras, dual-channel simultaneous acquisition. Excitation 488 and 561 nm; emission 498 nm (CaL520) and 571 nm (reference). Lateral sampling 0.183 µm px^−1^ over 1920 × 1920 px (351 × 351 µm), single plane. Frame interval 0.5293 s (1.89 Hz), except three recordings at 0.2996 s (3.34 Hz), the two vehicle recordings and one no-stimulus control; recording durations 11–202 s.

#### Analysis

Stacks were registered to their first frame by rigid translation estimated on the reference channel, which carries no indicator. Active cells were marked by eye, an automated variance-based detector having been evaluated and rejected at 61% recall and 13% precision, and were segmented as the half-maximum connected component of the background-subtracted time mean. Traces are ΔF/F against each cell’s own 20th percentile with a running baseline removed; a transient was counted when its prominence exceeded 4σ with a rise under 1.6 s and decay to 40% of its prominence within 8 s, detected identically in the reference channel. Frames in which a cell’s local contrast fell below half its own 99th percentile were excluded as out of focus, and frames in which more than 8% of structure regions stepped by more than 3 robust SD were vetoed as z excursions, along with events within one frame of them, in both channels. Cells whose only event was a single transient in the first 10 s were discarded, that being within the no-dye false-positive range, as were cells left with no events (15 of 75 cells in total). Quantities are aggregated cells to recording to organ to animal, so that an organ imaged twice contributes once; condition values are the median over animal means. Comparisons are two-sided Mann–Whitney tests with the design floor stated alongside, and the design is a survey, so every *P* value is descriptive. Manual marks had a median radius of 6.59 µm (range 1.83–10.89) and were used for duplicate merging and for a 4 µm seed search only, never as cell boundaries; the noise annulus was the segmented mask dilated from 3 µm to 10 µm. Dominant frequency was taken as the peak of a Welch spectrum (Hann window, segments of min(n, 128) samples, 50% overlap, mean removed) within 0.1–0.9 Hz, the upper bound set by the Nyquist limit at the predominant 1.89 Hz sampling, and transient rates were measured in the first 30 s of each recording. The Clark–Evans index is reported without edge correction and is therefore biased in small patches, and no separate photobleaching correction was applied: the running baseline removes slow drift, and the within-recording decline in activity is reported rather than corrected. Analysis used Python 3.12.15 with NumPy 2.5.4, SciPy 1.18.1, scikit-image 0.26.0 and pandas 2.2.3.

### Statistics

Averaging follows organ ⍰ animal ⍰ species, and *n* denotes animals unless organs are stated. Tests are two-sided: Fisher’s exact test for proportions, the Mann–Whitney *U* test for unpaired distributions, and the Wilcoxon signed-rank test for paired organ-versus-body-surface comparisons. Medians and ranges are reported rather than means and standard deviations, because several quantities span more than two orders of magnitude. The dataset comprises 24 organs from 15 animals. Species comparisons are restricted to ages at which all species were recorded (≤72 h); the age comparison is carried by *E. berryi*.

### Software and data

Analysis used Python 3.12 with NumPy 2.5, SciPy 1.18, pandas 2.2, scikit-image 0.26, tifffile and Matplotlib 3.11; PIV in PIVlab 3.14 under MATLAB R2025a; *D. pealeii* particle tracking in TrackMate 8.1.3 under Fiji; region drawing and projections in ImageJ 1.53c.

## Acknowledgements

We thank S. Colin and J. Costello, Spencer Nyholm, Margaret McFall-Ngai, Rachel Whitaker and the Marine Biological Laboratory (MBL) Microbial Diversity course, the MBL Image Analysis Lab and Central Microscopy Facility (especially L. Kerr), and the Cephalopod Breeding Program at the MBL Marine Resources Center. The fluorescent *Vibrio fischeri* ES114 strains were a gift from the Nyholm laboratory, the *Vibrio alginolyticus* strain from Rachel Whitaker and the MBL Microbial Diversity course, and the EsPGRP4 antiserum from the McFall-Ngai laboratory. This work was supported by an MBL Whitman Fellowship (J.C.N.).

## Figure legends

**Figure S1.**
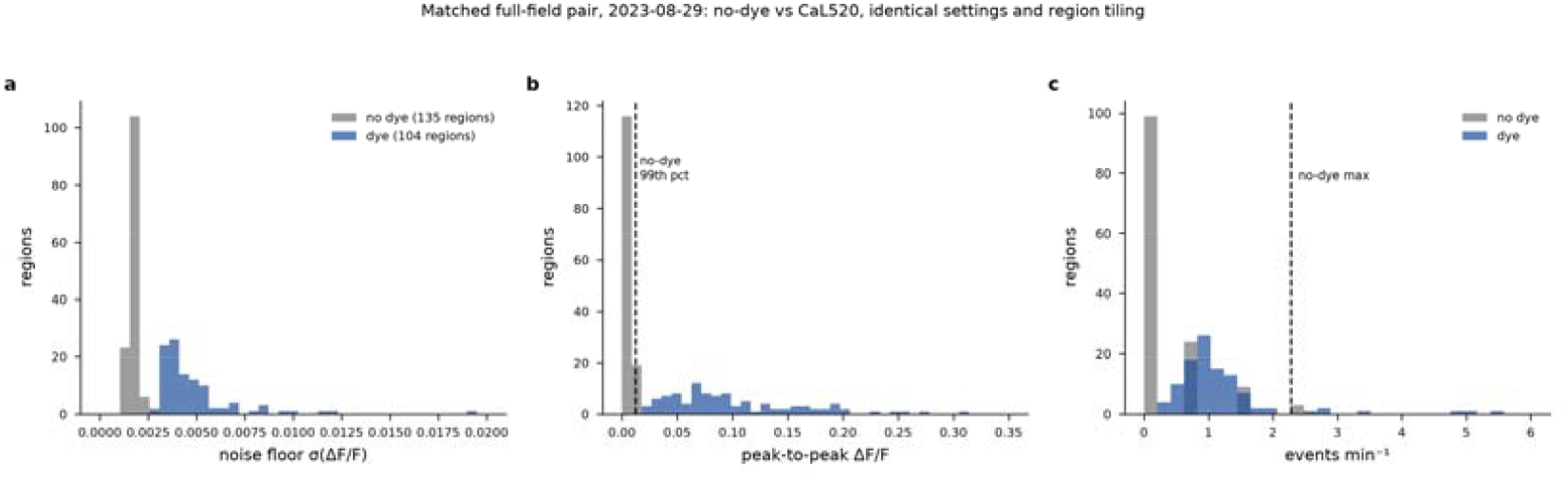
No-dye controls bound the detector’s artefact floor. Matched full-field pair from one cohort: one animal loaded with CaL520 and one receiving the DMSO vehicle without indicator, imaged at identical settings and tiled identically with cell-sized regions. (**a**) Noise floor σ(ΔF/F) per region. (**b**) Peak-to-peak ΔF/F per region; dashed line, the no-dye 99th percentile. (**c**) Detected events per minute per region; dashed line, the highest rate any no-dye region reaches (2.28 min^−1^). All three distributions separate at *P* = 5 × 10^−40^ (Mann–Whitney, 135 no-dye and 104 dye regions).

**Figure S2.**
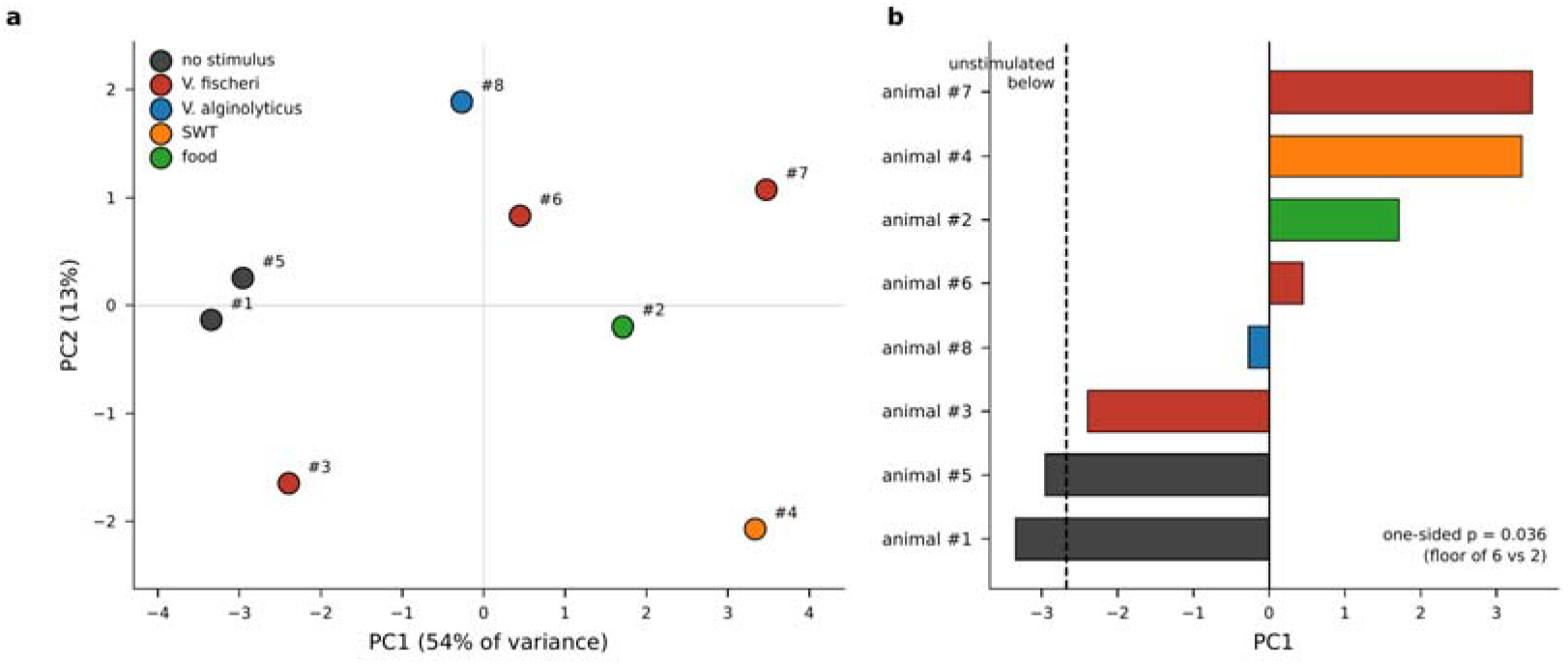
Multivariate structure separates stimulus vs non-stimulus conditions. Each animal is summarised as its median over twelve per-cell metrics (transient rate, event count, cell area and equivalent diameter, eccentricity, dominant frequency, periodic power ratio, indicator amplitude, indicator-to-reference amplitude ratio, signal-to-noise, inter-event-interval CV, and whole-recording rate), z-scored across animals, and entered into a principal component analysis; numbers identify the animals of Figure 4. (**a**) The first two components. PC1 carries 54% of the between-animal variance and is loaded near-equally on signal-to-noise (0.37), equivalent diameter (0.36), cell area (0.35), transient rate (0.34) and amplitude ratio (0.34), so it is a general axis of more and larger active cells rather than any single measurement; PC2 carries 13% and separates nothing. (**b**) The same animals ranked on PC1 alone. The two unstimulated animals are the lowest two of eight, a complete separation from the six stimulated animals (dashed line); Mann–Whitney p = 0.036 one-sided and 0.071 two-sided, both the smallest values attainable with six animals against two. The analysis is unsupervised, so the component is defined without reference to the condition labels. 60 cells, 10 organs, 8 animal*s*.

**Figure S3.**
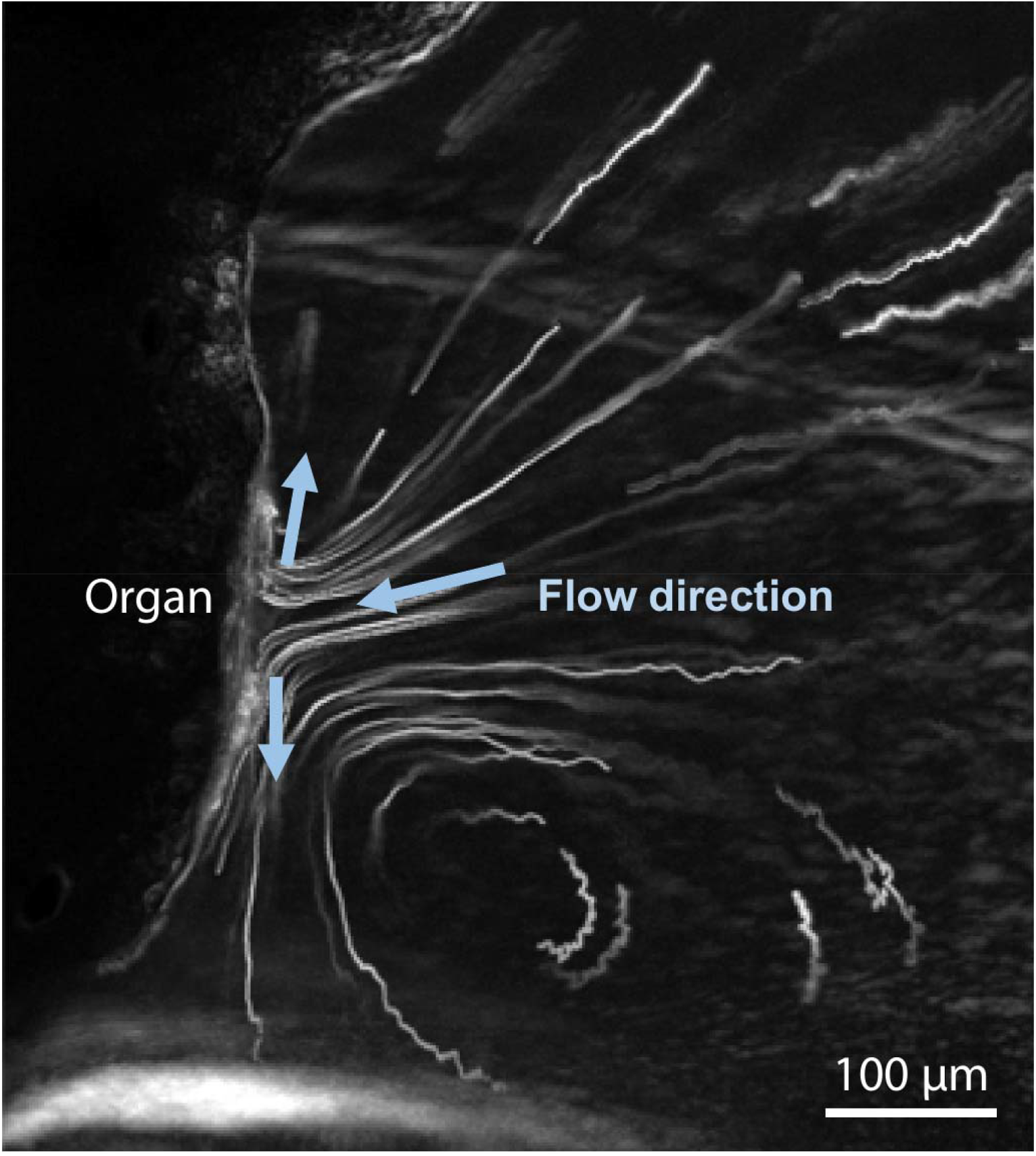
Flow pattern is consistent with phase reversal of ciliary beat. Flow tracer trajectories illustrate advection of the flow to the ciliated surface where the flow is separates into opposite directions, consistent with the observation of phase reversal of ciliary beat near the center of the organ.

